# How sensitive is functional connectivity to electrode resampling on intracranial EEG? Implications for personalized network models in drug-resistant epilepsy

**DOI:** 10.1101/696476

**Authors:** Erin C. Conrad, John M. Bernabei, Lohith G. Kini, Preya Shah, Fadi Mikhail, Ammar Kheder, Russell T. Shinohara, Kathryn A. Davis, Danielle S. Bassett, Brian Litt

**Author notes:** These authors contributed equally (Erin C. Conrad, John M. Bernabei). Corresponding Author: Erin C. Conrad, MD, Department of Neurology, Hospital of the University of Pennsylvania, 3400 Spruce Street, Philadelphia, PA 19104.

## Abstract

Focal epilepsy is a clinical condition arising from disordered brain networks. Network models hold promise to map these networks, localize seizure generators, and inform targeted interventions to control seizures. However, incomplete sampling of epileptic brain due to sparse placement of intracranial electrodes may profoundly affect model results. In this study, we evaluate the robustness of several published network measures applied to intracranial electrode recordings and propose an algorithm, using network resampling, to determine confidence in model results. We retrospectively subsampled intracranial EEG data from 28 patients who were implanted with grid, strip, and depth electrodes during evaluation for epilepsy surgery. We recalculated global and local network metrics after both randomly and systematically resampling subsets of intracranial EEG electrode contacts. We found that sensitivity to incomplete sampling varied significantly across network metrics, and that this sensitivity was independent of the distance of removed contacts from the seizure onset zone. We present an algorithm, using random resampling, to compute patient-specific confidence intervals for network localizations on both global and nodal network statistics. Our findings highlight the difference in robustness between commonly used network metrics and provide tools to assess confidence in intracranial network localization. We present these techniques as an important step toward assessing the accuracy of intracranial electrode implants and translating personalized network models of seizures into rigorous, quantitative approaches to invasive therapy.

## 1. Introduction

Epilepsy is a significant cause of disability worldwide, particularly among the one-third of patients whose seizures cannot be controlled by medications (Kwan, Schachter, and Brodie 2011; Wiebe et al. 1999). While these patients may benefit from surgery or implanted devices,many continue to experience seizures after invasive therapies (Engel 1996; Wiebe et al. 2001; Englot, Birk, and Chang 2017; Noe et al. 2013). One reason for this persistence of seizure activity may be the difficulty in localizing seizure-generating brain regions, the drivers of complex epileptic brain dynamics.

Clinicians and scientists now agree that epilepsy is in part a disease of brain networks (Kramer and Cash 2012). Driven by clinical observations, scientists have applied formal models from network theory to better understand seizure dynamics and target therapy (Bassett, Zurn, and Gold 2018). In these models, the brain is discretized into regions represented by network nodes, while network edges are used to represent their structural or functional connectivity. Network theory applied to epilepsy employs a wide variety of metrics to understand seizure generation and control, including node strength (Proix et al. 2017), betweenness centrality (Wilke, Worrell, and He 2011), clustering coefficient (Liao et al. 2010), and control centrality (Khambhati et al. 2016), as well as global metrics including global efficiency (Pedersen et al. 2015), synchronizability (Khambhati et al. 2016), and transitivity (Paldino et al. 2017). Collectively, these network measures have been used to predict neuronal firing as seizures begin and spread, track seizure progression, identify the seizure onset zone, and predict surgical outcome (Burns et al. 2014; Ponten, Bartolomei, and Stam 2007; Fletcher and Wennekers 2018; Wilke, Worrell, and He 2011; Panzica et al. 2013; Sinha et al. 2016).

When using invasive sensors such as intracranial EEG (iEEG) to estimate functional connectivity, sampling from the full brain is impossible, and the network measures available for modeling depend on the location and number of electrodes implanted. In fields outside of epilepsy, missing data is known to affect the results of network analyses (Albert, Jeong, and Barabási 2000; Albert, Albert, and Nakarado 2004; Guimerà and Sales-Pardo 2009; Lü and Zhou 2011). The effect of missing data on network models and clinical care in epilepsy has not been rigorously explored. While network models have potential to add rigor to clinical decision making, their application may be limited by uncertainty in estimated network metrics and the unknown interaction between that uncertainty and sparse brain sampling. In this study we seek to rigorously assess the extent to which different network metrics are sensitive to intracranial electrode sampling. This computational work is a vital first step to deploying network models as an adjunct to clinical decision-making.

## 2. Materials and Methods

### 2.1 Summary

We use a high-quality dataset that has been included in multiple network studies in epilepsy (Khambhati et al. 2016; Sinha et al. 2016; Khambhati et al. 2015) and is publicly available at www.IEEG.org. We randomly eliminate nodes from functional networks to simulate the uncertainty consequent to variable sampling of brain regions by iEEG and to determine the reliability of different network metrics within and across patients. Based upon the assumption that the main drivers of epilepsy network behavior might localize to an epileptogenic region, we ask to what extent electrode contacts far away from the seizure onset zone impact the estimated values of various network metrics. We then randomly remove nodes by jackknife resampling in order to derive patient-specific estimates of confidence in network statistics.

### 2.2 Patient selection, intracranial EEG recording, and electrode localization

All patients gave written informed consent in accordance with the Institutional Review Board of the Hospital of the University of Pennsylvania (HUP) and the Mayo Clinic in Rochester. Furthermore, all patients consented to publishing their full length iEEG recordings on the public web portal IEEG.org (Wagenaar et al. 2013). This study was performed in accordance with the Declaration of Helsinki.

A total of 28 patients with drug-resistant epilepsy underwent iEEG recording during presurgical evaluation at HUP or the Mayo Clinic. Electrode configurations (Ad Tech Medical Instruments, Racine, WI) consisted of linear cortical strips and two-dimensional cortical grid arrays (2.3 mm diameter with 10 mm inter-contact spacing), and linear depth electrodes (1.1 mm diameter with 10 mm inter-contact spacing). EEG signals were recorded at a sampling frequency of 512 Hz at HUP and 500 Hz at Mayo Clinic. All electrode and EEG recording systems were FDA approved and are commercially available.

Each patient underwent MPRAGE T1-weighted magnetic resonance imaging (MRI) on a 3T Siemens Magnetom Trio scanner (Siemens, Erlangen, Germany) prior to electrode implantation, and they also underwent spiral CT imaging (Siemens, Erlangen, Germany) after electrode implantation. We cross-referenced the CT images with a surgical cartoon map to localize electrode coordinates (Wu et al. 2011). To segment the resection zone, we registered the pre-implant MRI to post-resection imaging and the post-implant CT using the Advanced Normalization Toolkit (ANTs) (Avants et al., 2011). We utilized a random forest classifier with ITK-SNAP to semi-automatically estimate the resection zone and identify electrodes overlying resected cortex (Yushkevich et al., 2006).

Seizures were identified clinically and confirmed in a clinical case conference discussion. Board-certified epileptologists (Fadi Mikhail, Ammar Kheder, Kathryn Davis, and Brian Litt) then reviewed the seizures and identified the earliest electrographic change (EEC) (Litt et al. 2001) and the electrode contacts of seizure onset for each seizure. We performed our primary analysis on the first seizure identified for each patient. For patients with more than one seizure (N = 26), we also performed the analysis on the second seizure to assess the sensitivity of our results to the choice of seizure. One patient (HUP111) had two separate electrode implantations, and we analyzed both implantations separately.

### 2.3 Calculating Functional Networks

We examined one-second time windows at each of the following time periods: 10 seconds prior to the EEC, 5 seconds prior to the EEC, at the EEC, 5 seconds after the EEC, and 10 seconds after the EEC. We performed our primary analysis on the time period at the EEC, which we anticipated to be the period most important for understanding peri-ictal networks. We then repeated the analysis for each other time window in order to assess the sensitivity of our results to the choice of time period.

A common average reference was applied to iEEG signals to remove common sources of noise. Data were filtered using an elliptic bandpass filter with cutoff frequencies of 5 Hz and 115 Hz, as well as a 60 Hz notch filter to remove power line noise. Signals were pre-whitened using a continuous autoregressive model to account for slow dynamics and to accentuate higher frequencies known to be involved in seizure dynamics. For each one-second window, we constructed functional networks in which iEEG electrode contacts represented network nodes. Edges were weighted by multitaper coherence, which estimates the correlation between two electrode contact signals in the frequency domain (Mitra and Pesaran 1999). We calculated coherence in the high gamma frequency band (95-105 Hz), which we chose due to its importance in seizure propagation and spread (Khambhati et al. 2016). We also repeated the analysis in beta (15 - 25 Hz) to assess the sensitivity of our results to the choice of frequency band, and in acknowledgment of the fact that the beta frequency is thought to be important in epileptic networks as well (Bettus et al. 2011). This separation of the data resulted in an adjacency matrix for each frequency band representing a network with undirected, weighted edges for each patient, where each row and each column represented an electrode contact, and each matrix element represented the signal coherence between the two contacts.

### 2.4 Network metrics

For each functional network, we calculated several global and nodal network metrics, chosen because of their importance in graph theory and their use in recent epilepsy publications as described above. The global metrics were synchronizability, global efficiency, and transitivity. The nodal metrics were node strength, control centrality, clustering coefficient, eigenvector centrality, and betweenness centrality. The methods for calculating these metrics have been previously described, and we briefly summarize each below. We specifically describe their calculations for an undirected, weighted network. We calculated each using the Brain Connectivity Toolbox (Rubinov and Sporns 2010), or using custom code for synchronizability and control centrality.

Node strength represents the total strength of connections involving a particular node (Fornito, Zalesky, and Bullmore 2016), and can be defined as

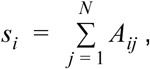

in which *s*_*i*_ is the strength of node *i, A*_*ij*_ is the adjacency matrix element containing the edge weight between node *j* and node *i*, and *N* is the number of nodes. A high node strength implies that the total weight of its connected edges is large. Eigenvector centrality is a similar nodal measure that weights individual node influence by the relative influence of each of its connected nodes (Newman 2008; Fletcher and Wennekers 2018). It is specifically defined as λ = *Ax* where *x* is the vector containing the eigenvector centrality of each node, *A* is the adjacency matrix, and *λ* is the largest eigenvalue of the matrix (which results in non-negative eigenvector centralities). A high eigenvector centrality implies that a node is strongly connected to nodes that themselves are highly connected.

Global efficiency is a global measure that is thought to represent how easily information travels throughout the network (Latora and Marchiori 2001). It is defined as

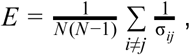

where *E* is global efficiency, *N* is the number of nodes, and σ_*ij*_ is the shortest weighted path length between node *i* and node *j*, for example estimated using Dijkstra’s algorithm (Dijkstra 1959). A high global efficiency is thought to reflect a greater ease of information transfer throughout the network (Bassett et al. 2009). Path lengths were weighted by the inverse of the values of the adjacency matrix, to reflect the fact that information is thought to be transferred more readily along stronger edges (Opsahl, Agneessens, and Skvoretz 2010). Betweenness centrality is a related nodal metric that measures the fraction of all shortest paths in the network that pass through a given node (Freeman 1977). It is defined as

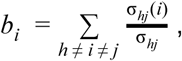

where *b*_*i*_ is the betweenness centrality of node *i*, σ_*hj*_*(i)* is the number of shortest paths from node *h* to node *j* that pass through node *i*, and σ_*hj*_ is the total weighted path length between node *h* and node *j*. A high betweenness centrality suggests that the node acts as a central node in the shortest paths between many other nodes. The path lengths were weighted by the inverse of the values of the adjacency matrix as described above.

Synchronizability is a global metric that quantifies the stability of the fully synchronous network state (Schindler et al. 2008; Boccaletti et al. 2006) and has been shown to predict seizure generalization (Khambhati et al., 2016). It is calculated by first computing the weighted Laplacian *L* = *D* - *A* as the difference between the node strength matrix *D* and the adjacency matrix *A*. Synchronizability is then obtained by the equation 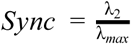, where *Sync* is synchronizability, *λ*_*2*_ is the second smallest eigenvalue of the Laplacian, and *λ*_*max*_ is the largest eigenvalue. Greater synchronizability reflects a smaller spread between eigenvalues, which suggests greater ease for a network to synchronize its dynamics. Control centrality is a corresponding local metric that measures the effect of each node on synchronizability. It is defined as 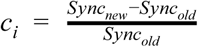, where *c*_i_ is the control centrality of node *i, Sync*_*old*_ is the original synchronizability, and *Sync*_*new*_ is the synchronizability of the network with the node(s) removed (Khambhati et al. 2016). Negative control centrality nodes are synchronizing, whereas positive control centrality nodes are desynchronizing.

Transitivity is another global measure that represents the degree to which nodes in a graph tend to cluster together (Holland and Leinhardt 1971; Watts and Strogatz 1998; Opsahl and Panzarasa 2009). It is defined as

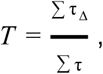

where *T* is transitivity, ∑τ _Δ_ is the sum of the weights of closed triplets,and ∑τ is the sum of the weights of all triplets. A triplet is defined as a set of three nodes connected by either two or three edges. A closed triplet, more specifically, is a set of three nodes connected by three edges. Higher transitivity implies that nodes tend to cluster together into exclusive groups. The clustering coefficient is the nodal extension of transitivity that measures the amount of interaction between local triplets (Barrat, Barthélemy, and Vespignani 2007). It is calculated by 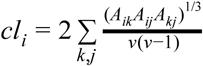 in which *A* is the adjacency matrix edge weight and *v* is the number of neighbors. Higher clustering coefficients reflect greater clustering of the node into tight groups.

### 2.5 Network resampling

To determine the sensitivity of network metrics to spatial sampling, we randomly identified electrode contacts for removal in each patient. We removed the rows and columns corresponding to these electrode contacts from the adjacency matrix representing the network. We recalculated each of the network metrics above. We performed this analysis removing 20%, 40%, 60%, and 80% of randomly selected electrode contacts. We repeated this process 1,000 times for each removal percentage to obtain a distribution of new metrics in the randomly resampled network (Fig. 1).

**Figure 1:**
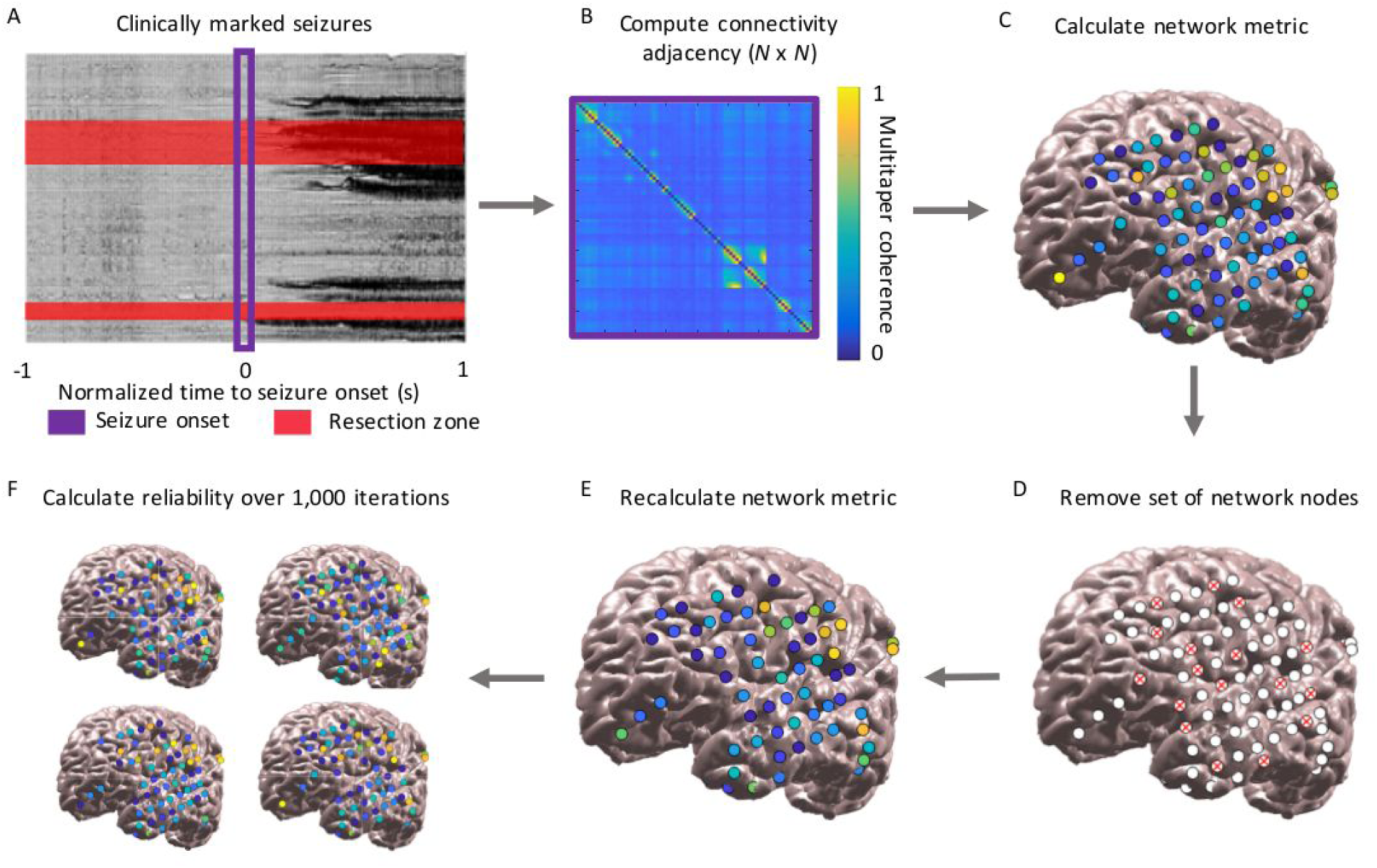
Network generation and resampling methods. A: Seizure onset times were marked by a board-certified epileptologist. Resected brain areas corresponding to EEG channels were determined through a semi-automated imaging technique and confirmed by a board-certified neuroradiologist. B: Multitaper coherence of a 1-second interval of EEG signal at seizure onset was used to create a functional adjacency matrix. C: Network metrics were calculated using the adjacency matrix with all nodes included. D: A subset of nodes were removed to simulate the effect of leaving out electrodes. E: Network metrics were recalculated from the resampled network. F: This process was repeated over 1,000 iterations and the reliability of each metric was quantified.

To determine the stability of each network metric to resampling, we calculated the *reliability* for each removal percentage (Davidshofer and Murphy 2005). Reliability is defined as 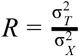, where 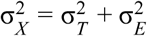, and *σ*^*2*^_*T*_ is the variance of the true scores, *σ*^*2*^ _*E*_ is the variance of th error, and *σ*^*2*^_*X*_ is the total variance. We defined the variance of the error to be the variance of the resampled metric across the 1,000 random resamples, averaged across electrode contacts in the case of nodal metrics. For nodal metrics, we defined the variance in the true scores to be the variance of the resampled metric across electrode contacts, averaged over all permutations. In the case of global metrics, we defined the variance in the true scores to be the variance in the resampled metric across patients, averaged over all permutations. Reliability is constrained to be between 0 and 1, where 1 indicates that no variance is due to random resampling, 0 indicates that all variance is due to random resampling, and 0.5 indicates that the variance due to random resampling equals the variance of the true metric. To determine whether some metrics were more robust to resampling than others, we compared the metric reliability across all patients for the 20% removal percentage using separate Friedman tests, one for global metrics and one for nodal metrics (α = 0.05) (Friedman 1937). In the case of significant Friedman test results, we performed *post hoc* Dunn-Sidak’s multiple comparisons tests to identify significant differences between individual metrics (Dunn 1964; Šidák 1967). We also determined the reliability of metrics for removal percentages other than 20%, which we report in our supplemental data. We repeated this analysis for beta band coherence, alternate times relative to the EEC, and removal of contiguous rather than random electrode contacts, which we also report in our supplemental data.

We then determined whether there was a relationship between the network reliability and the number of electrode contacts comprising the original network. We obtained the reliability for each patient and each nodal and global metric at the 20% removal percentage of random electrode contacts, using the EEC time period and gamma band coherence. For each metric, we correlated the reliability with the original number of electrode contacts in the patient’s network using Spearman rank correlation. We performed Bonferroni correction as we were testing eight network metrics, yielding an *α* of 0.05/8 = 0.00625.

We next hypothesized that ictal network metrics may be more affected by removing electrode contacts near the seizure onset zone, as these contacts may have a stronger influence on epileptic networks. To test this, we again resampled the network, this time systematically removing each electrode contact and its *N*-1 nearest neighbors, where *N* was equal to 20% of the total number of contacts in the network (we also calculated it for other removal percentages and report these results in our supplemental data). We recalculated each of the global and nodal metrics in this systematically resampled network. We obtained a measure of agreement between the original metric and the new metric in the resampled network. For nodal metrics, the agreement measure *a* was defined as the Spearman’s rank correlation coefficient across electrode contacts between the original and resampled metric. For global metrics, the agreement measure was defined as the negative of the absolute value of the relative difference between the two metrics

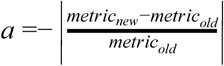

The global agreement *a* was equal to 0 when there was perfect agreement between the new and original metric, and was increasingly negative with larger absolute differences.

To test whether there was larger metric agreement when the removed electrode contacts were further from the seizure onset zone, we obtained the Spearman’s rank correlation coefficient between the agreement measure *a* with the distance between the centroid of the removed electrode contacts and the centroid of the seizure onset zone. We obtained the Fisher’s transformation of the rank coefficient for each patient, which is equal to the inverse hyperbolic tangent of the rank coefficient, in order to transform the coefficients to a variable whose distribution is approximately normal (Fisher 1915). We aggregated these transformed rank coefficients across patients and performed a two-sided one-sample *t*-test to determine if the mean coefficient was significantly different from zero. We performed this test for each of the global and nodal metrics. We performed Bonferroni correction as we were testing eight network metrics, yielding an *α* of 0.05/8 = 0.00625.

We next utilized a jackknife resampling method to generate patient-specific estimates in the confidence of the results of network analyses. Jackknife estimation is a method of sampling without replacement to derive statistical estimates (Quenouille 1949, 1956; Tukey 1958). Our goal was to determine how much a network result would be expected to change if a small number of electrode contacts had not been present. We randomly removed 20% of electrode contacts, recalculated the network statistic of interest for the random resample, and repeated this process for 1,000 iterations. We chose a 20% removal percentage for this analysis to simulate minor variability in electrode implantation strategy. For each of the nodal metrics, we performed this analysis to determine the electrode contacts accounting for 95% of occurrences of the maximal value of the metric (minimal value for control centrality). We called this set of contacts the 95% jackknife confidence contact set. We also identified the 95% confidence contact set of the minimum *regional control centrality*, defined as the locations of an electrode contact and its *N*-1 nearest neighbors, where *N* equals the number of resected electrode contacts that produces the largest negative change in synchronizability when removed. Regional control centrality attempts to identify a region of a defined size - rather than a single electrode contact - with the largest control centrality, and thus a potential site for resection. For global metrics, we performed this method to obtain the 95% jackknife confidence interval for the value of the metric for a given patient, which was the interval containing 95% of all values obtained with jackknife resampling. The runtime for the jackknife resampling algorithm (1,000 iterations) for all metrics at a single time and frequency band was approximately ten minutes per patient when performed in MATLAB R2018a on an Intel® Xeon® processor (CPU E5-2698 v3 @ 2.30GHz).

### 2.6 Statistical analysis

All analyses were performed on MATLAB R2018a (The Mathworks, Natick). Specific analyses are discussed in Section 2.4.

### 2.7 Data and code availability

All EEG records and annotations are available on The International Epilepsy Electrophysiology Portal (https://www.ieeg.org/) (Wagenaar et al. 2013; Kini, Davis, and Wagenaar 2016). All code is available on GitHub (https://github.com/erinconrad/network-subsets).

## 3. Results

### 3.2 Patient and electrode information

Patients had a variety of clinical histories, electrode configurations, pathologies, and clinical outcomes (Supplemental Table 1). There were 28 patients (13 women), one of whom had two temporally distinct implantations, which were separately analyzed. The average age at implantation was 33.9 years (range 5-57). The mean number of electrode contacts was 77 (range 16-118). The mean number of seizures was 6.8 (range 1-36). The median International League Against Epilepsy (ILAE) outcome score at 2 years was 2 (range 1-5).

### 3.2 Stability of metrics to random resampling

For all network measures, reliability to resampling decreased as more electrode contacts were removed. The stability of network measures to resampling varied across patients (Fig. 2). The mean reliability was *R* = 0.92 for synchronizability, *R* = 0.98 for global efficiency, and *R* = 0.98 for transitivity, averaged over all patients when a random sample of 20% of electrode contacts was removed. The reliability was significantly different between global metrics (Friedman test: *χ*^2^_2_= 36.5, *p* < 0.001). Synchronizability was significantly less reliable than either global efficiency or transitivity (*post hoc* Dunn-Sidak’s multiple comparison test: *t* = −3.02, *p* = 0.008 compared to global efficiency and *t* = −6.04, *p* < 0.001 compared to transitivity). Global efficiency was also significantly less reliable than transitivity (*t* = −3.02, *p* = 0.008). The reliability for global efficiency was slightly lower than that for transitivity for 26 out of 29 patient implantations. However, for two implantations the reliability for global efficiency was substantially larger, explaining why global efficiency and transitivity have similar means despite global efficiency having significantly lower reliability by ordinal statistics.

**Figure 2:**
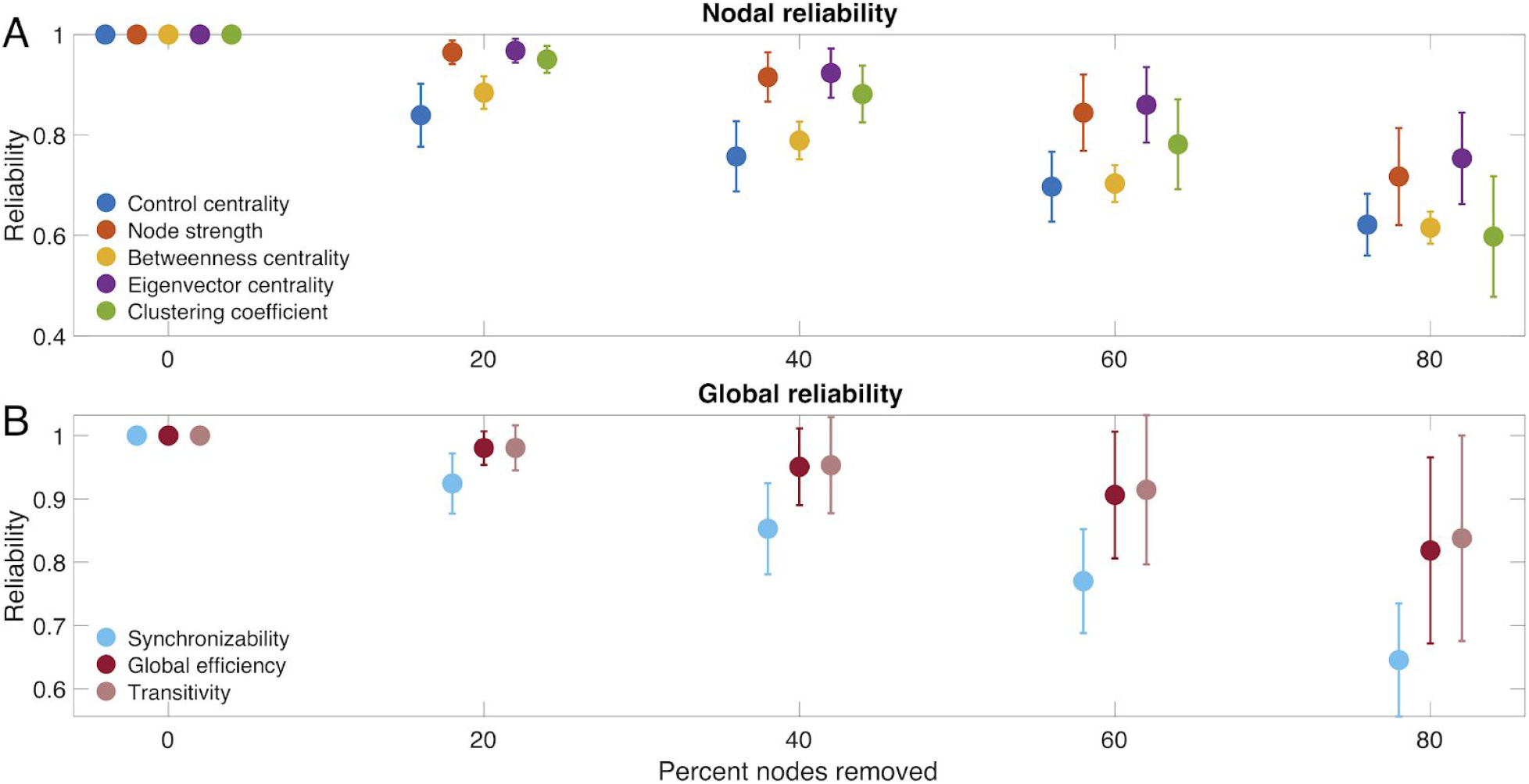
Reliability of network metrics to incomplete sampling. A: Reliability of nodal measures, averaged across all patients, when different percentages of electrodes were removed. B: Reliability of global measures, averaged across all patients, when different percentages of electrodes were removed. Error bars show the standard deviation of the reliability across patients. All data shown is for the EEC of first seizure, high gamma coherence, and random electrode removal. For all measures, reliability decreased as more electrodes were removed. Patients are heterogeneous in the reliability of their network measures, and certain network measures exhibit higher reliability than others.

For nodal measures, the mean reliability was *R* = 0.84 for control centrality, *R* = 0.96 for node strength, *R* = 0.88 for betweenness centrality, *R* = 0.97 for eigenvector centrality, and *R* = 0.95 for clustering coefficient, averaged over all patients when 20% of electrode contacts were randomly removed. The reliability differed significantly between nodal measures (Friedman test: *χ*^2^_4_ = 107.9 *p* < 0.001). Control centrality was less reliable than node strength (*post hoc* Dunn-Sidak test: *t* = −6.73, *p* < 0.001), eigenvector centrality (*t* = −8.64, *p* < 0.001), and clustering coefficient (*t* = −3.99, *p* = 0.001) but was not significantly different from betweenness centrality (*t* = −1.00, *p* = 0.979). Node strength was also significantly more reliable than betweenness centrality (*t* = 5.73, *p* < 0.001) but did not differ significantly from eigenvector centrality (*t* = −1.91, *p* = 0.439) or clustering coefficient (*t* = 2.74, *p* = 0.060). Betweenness centrality was significantly less reliable than both eigenvector centrality (*t* = −7.64, *p* < 0.001) and clustering coefficient (*t* = −2.99, *p* = 0.028). Eigenvector centrality was more reliable than clustering coefficient (*t* = 4.65, *p* < 0.001).

When we examined the time periods 10 seconds before, 5 seconds before, 5 seconds after, and 10 seconds after the EEC (as opposed to the second at the EEC), synchronizability continued to have the lowest reliability of the global metrics. Control centrality continued to have the lowest reliability of the nodal metrics, and eigenvector centrality and node strength continued to have the highest reliabilities. The pattern remained when we examined beta frequency coherence rather than high gamma frequency coherence, when we removed contiguous as opposed to random sets of electrode contacts, and when we examined the second rather than the first seizure (Supplemental Table 2). When we instead removed 40% or 60% of electrode contacts, control centrality and synchronizability continued to have the lowest reliability of nodal and global metrics, respectively, and node strength and eigenvector centrality continued to have the highest nodal metric reliabilities. When we removed 80% of electrode contacts, clustering coefficient instead demonstrated the lowest nodal metric reliability, and otherwise the pattern was unchanged (Supplemental Table 3).

We examined the relationship between robustness to electrode contact resampling and the number of electrode contacts in the original network. Amongst global measures, there was a significant positive relationship between reliability and number of contacts for synchronizability (Spearman rank correlation: *r*_27_ = 0.68, *p* < 0.001), but not for global efficiency (*r*_27_ = 0.20, *p* = 0.299) or transitivity (*r*_27_ = 0.22, *p* = 0.260). Amongst nodal measures, clustering coefficient (*r*_27_ = 0.53, *p* = 0.003) demonstrated a significant positive relationship, and relationships for control centrality (*r*_27_ = −0.12, *p* = 0.551), node strength (*r*_27_ = 0.45, *p* = 0.015), betweenness centrality (*r*_27_ = 0.38, *p* = 0.043), and eigenvector centrality (*r*_27_ = 0.44, *p* = 0.016) were non-significant (*α* = 0.00625, Bonferroni correction for eight measures). This pattern of findings suggests that, at least for synchronizability and clustering coefficient, patients with more electrode contacts implanted were less vulnerable to incomplete spatial sampling.

### 3.3 Influence of distant electrode contacts on clinical region of interest

There was no significant association between metric agreement and distance of the removed electrode contacts from the seizure onset zone for any metric (one-sample two-sided *t*-test: control centrality, *t* = 0.80, *p* = 0.433; node strength, *t* = 1.25, *p* = 0.222; betweenness centrality, *t* = −0.95, *p* = 0.352; eigenvector centrality, *t* = 1.02, *p* = 0.318; clustering coefficient, *t* = 1.23, *p* = 0.230; synchronizability, *t* = −0.26, *p* = 0.793; global efficiency, *t* = 0.74, *p* = 0.469; transitivity, *t* = 0.37, *p* = 0.717). This pattern of findings implies that all metrics are equally sensitive to removing electrode contacts near versus distant from the seizure onset zone (Fig. 3). This result was invariant to the choice of peri-ictal time window, choice of frequency band, or choice of seizure (Supplemental Table 4) as well as to the choice of removal percentage (Supplemental Table 5).

**Figure 3:**
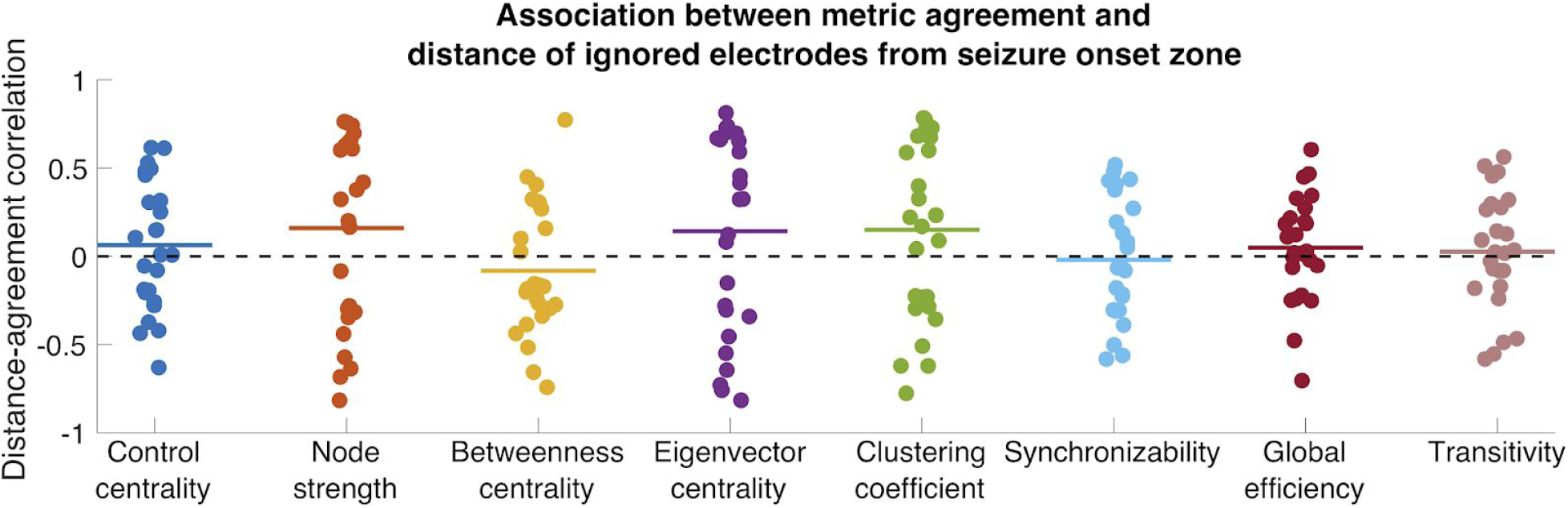
Association between metric agreement and distance of ignored electrodes from the seizure onset zone. Each nodal and global metric is shown. Each point represents the patient-specific Spearman rank correlation coefficient between the metric agreement and the distance of the ignored electrodes from the seizure onset zone. The metric agreement is defined for nodal metrics as the Spearman rank correlation coefficient between the original metric and the metric obtained from resampling, and for global metrics as the negative absolute value of the relative difference between the original and resampled metric. Horizontal lines show the average distance-agreement association across patients. No distance-agreement association was significantly different from zero (two-sided one-sample *t*-test, α = 0.05/8. Bonferroni correction), signifying that all metrics are equally vulnerable to incomplete sampling near versus distant from the seizure onset zone.

### 3.4 Jackknife confidence intervals

Both nodal and global metrics varied differentially across patients with respect to jackknife confidence intervals produced by resampling (Fig. 4A-C). The median and range for the number of electrode contacts accounting for 95% of all jackknife instances of the maximum nodal metric (minimum for control centrality) was 3 (range 2-5) for node strength, 4 (3-9) for betweenness centrality, 3 (2-5) for eigenvector centrality, 3 (2-5) for clustering coefficient, and 9 (4-22) for control centrality. The median number of electrode contacts accounting for 95% of all jackknife instances of the minimum regional control centrality (where the set of electrode contacts with minimum regional control centrality is the set, equal in number to the number of resected electrode contacts, that together produces the largest negative change in synchronizability when removed) was 48 (range 12-93). The median ratio between this number and the number of electrode contacts forming the true minimum regional control centrality set was 4.0 (range 1.6-16.0) (Fig. 4D). Regarding global metrics, the median width of the 95% jackknife confidence interval was 0.094 (range 0.045-0.192) for synchronizability, 0.016 (range 0.006-0.058) for global efficiency, and 0.012 (range 0.004-0.062) for transitivity (Fig. 4E). These results demonstrate the heterogeneity amongst patients in the level of confidence in estimated network statistics that can be revealed by the jackknife algorithm. The locations of electrode contacts with the maximum or minimum metric values, as well as the results of jackknife resampling, varied across time periods, choice of frequency band for coherence, and seizure (Supplemental Fig 1., Supplemental Table 6).

**Figure 4:**
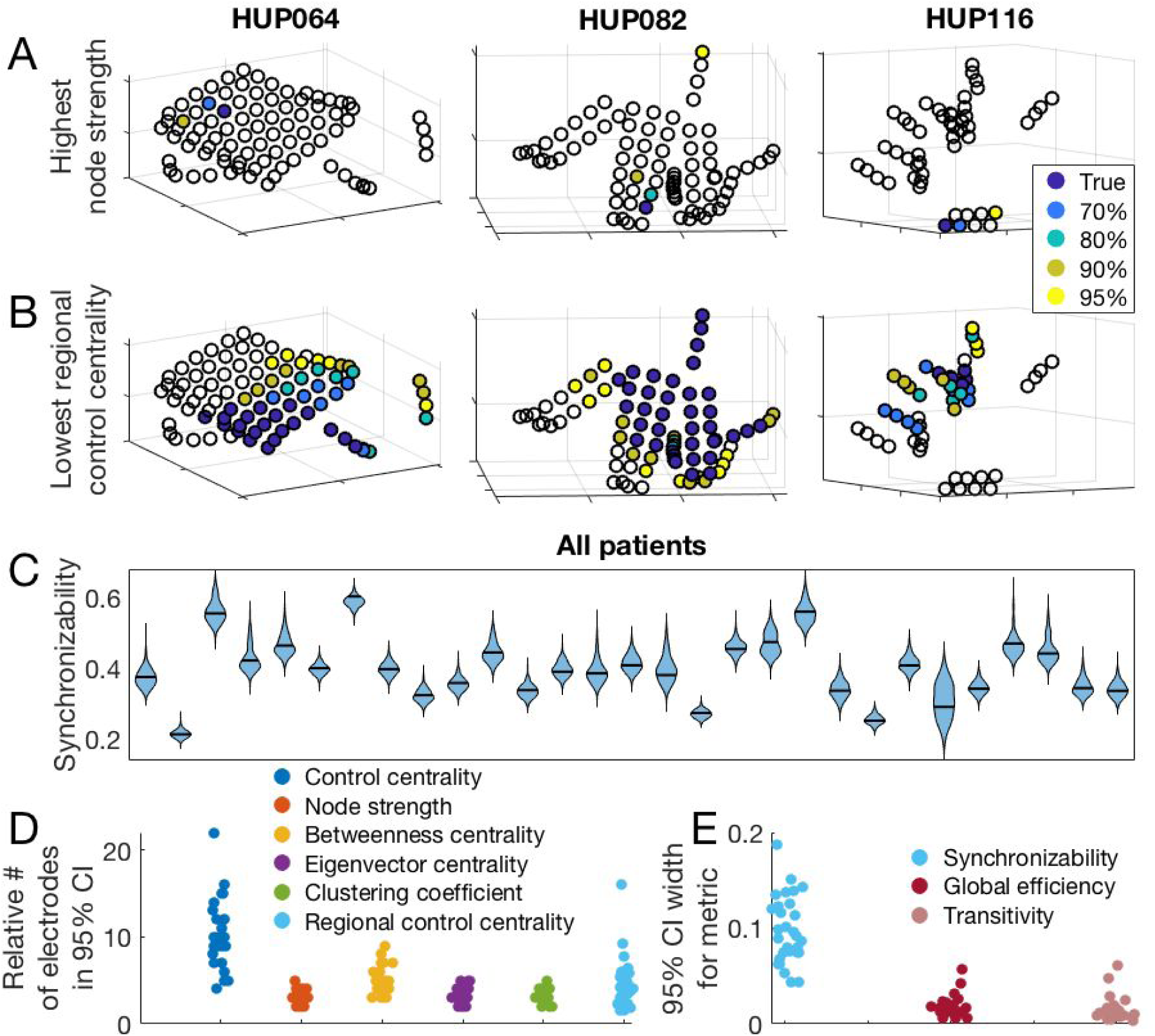
Jackknife resampling to estimate confidence regarding network metric values. A: The location of the electrode with the highest node strength, as well as the electrodes accounting for various percentages of highest node strength occurrences in 1,000 random jackknife resampling networks for three example patients. B: The location of the most synchronizing region (which is the region with the lowest regional control centrality), and the regions accounting for various percentages of the most synchronizing region occurrences in 1,000 random jackknife resampling networks for three patients. C: Patient-specific synchronizability distributions across resamples. Each separate violin represents a patient and shows the distribution of synchronizability values obtained across 1,000 random jackknife resamples. Horizontal black lines show the original value in the non-resampled network. D: Number of electrodes forming the 95% jackknife confidence electrode set for each nodal metric, for all patients. For each nodal metric, each dot shows the patient-specific number of electrodes accounting for 95% of all occurrences of the maximum (minimum for control centrality and regional control centrality) metric value in 1,000 random jackknife resampling networks. For regional control centrality, this number is divided by the number of electrodes forming the minimum regional control centrality in the original non-resampled network to obtain a ratio. E: Width of the 95% jackknife confidence interval for each global metric, for all patients. For each global metric, each dot shows the patient-specific width of the 95% jackknife confidence interval of the metric value across 1,000 random resamples. This figure demonstrates the variability in confidence of network theory results across patients that can be revealed by jackknife resampling.

## 4. Discussion

Handling missing data is a longstanding problem in science in general and is particularly problematic in network science, where a missing node may limit our understanding of the entire network (Guimerà and Sales-Pardo 2009). In social networks, missing data can dramatically alter network statistics (Kossinets 2006; Albert, Jeong, and Barabási 2000; Albert, Albert, and Nakarado 2004). In the field of neuroscience, Jalili demonstrated that global efficiency of scalp EEG-based functional networks in healthy individuals was highly sensitive to the removal of certain nodes (Jalili 2015). To our knowledge, this is the first study examining the reliability of network statistics in the epileptic brain and in iEEG data. We determined that network measures differ in robustness to spatial resampling, and that the sensitivity to sampling does not depend on the distance from the seizure onset zone. We also found that more extensive implants were more robust to sub-sampling. Finally, we developed and applied an algorithm using jackknife resampling of electrode contacts to estimate confidence in nodal and global statistics in patient brain networks.

### 4.1 Functional network metrics exhibit differential reliability under spatial resampling

Metric reliability for most network measures decreased with a greater degree of missing data, which has previously been reported in social networks (Smith and Moody 2013; Kossinets 2006). Among examined nodal metrics, node strength and eigenvector centrality were most reliable and control centrality was least reliable; among the global metrics we tested, transitivity was most reliable and synchronizability was least reliable. The difference in reliability across metrics reflects, in part, the underlying sensitivity of each metric to graph topology. Prior studies in social networks have also observed that node strength is more robust to resampling than betweenness centrality (Galaskiewicz 1991; Costenbader and Valente 2003; Smith and Moody 2013). The relative robustness of node strength and eigenvector centrality compared to other nodal measures suggests that metrics that depend only on immediate connections to the node of interest are less sensitive to resampling than metrics that incorporate multi-step connections. The preserved ordinality of network metric reliability across most patients, time scales, and frequency bands suggests that this result is generalizable. Clinically, applying network statistics that are more robust to spatial sampling may be preferable in cases in which the electrode coverage of important regions is uncertain. The ability of each metric to capture network behavior must be weighed against its spatial reliability if such personalized models are to be translated clinically.

Sensitivity to incomplete sampling depends somewhat on the number of electrode contacts forming the original implantation. Although synchronizability and clustering coefficient were the only global and nodal measures, respectively, to demonstrate a significant positive relationship between number of electrode contacts and metric reliability, all other measures except control centrality demonstrated non-significant positive relationships. This pattern of findings suggests that for most network measures, greater robustness can be achieved in part through more extensive electrode coverage. Mechanistically, this may imply that in larger networks, random subsampling is less likely to remove important hubs. Alternatively, perhaps information about missing nodes and edges can be inferred from the remaining components of the network, which is facilitated by a larger starting network. Clinically, this suggests that implanting a greater number of electrode contacts may increase our confidence in network statistics. This benefit would have to be weighed against the risks of more extensive coverage, including hemorrhage and infection (Mullin et al. 2016).

### 4.2 Metric sensitivity to incomplete sampling is independent of distance from the seizure onset zone

All metrics tested were equally sensitive to removing nodes in close or far proximity from the clinician-defined seizure onset zone. This may be because the seizure-generating network is a relatively small subset of the full peri-ictal network, and thus perturbation of the seizure-generating network has a small effect on network as a whole. Alternatively, functional importance to the seizure-generating network may be independent of spatial proximity to the seizure onset zone. This result is practically concerning in the case of metrics with lower reliability to spatial resampling, such as control centrality and synchronizability, because placement of electrodes in the network periphery away from clinical zones of interest is often variable across patients and across epilepsy centers. To increase the clinical confidence in the results of these network measures, the incomplete network may be supplemented using structural connectivity data and atlas-based approaches (Shah et al. 2018; Greicius et al. 2009; Liao et al. 2011; Fan et al. 2016; Betzel et al. 2017; Reddy et al. 2018). Network theory also proposes several methods of predicting missing links (Lü et al. 2015; Pan et al. 2016; Lü and Zhou 2011; Guimerà and Sales-Pardo 2009).

### 4.3 Jackknife network resampling generates confidence intervals for virtual resection

Prior work has used global and nodal network measures to stratify surgical candidates and to select nodes for resection (Tomlinson, Porter, and Marsh 2017; Sinha et al. 2016; Lopes et al. 2017; Shah et al. 2019). Here we provide a simple algorithm to augment these methods by determining patient-specific confidence in the robustness of estimated statistics to small perturbations in spatial sampling. Similar resampling-based approaches have been used in social networks (Duval, Christensen, and Spahiu 2010; Ohara et al. 2014; Lusseau, Whitehead, and Gero 2008), in gene expression data (de Matos Simoes and Emmert-Streib 2012), and in resting fMRI-data of healthy subjects (Cheng et al. 2017). The heterogeneity in confidence across patients may be used to stratify patients into those for which enough information of the epileptic network is captured to accurately use personalized network models and those for whom models are likely to be inaccurate due to implantation strategy. In this study we do not claim to identify the ideal network model among the many published studies. The jackknife resampling method may be easily applied to any of these network models.

### 4.4 Methodological limitations and future directions

Our method of network resampling can only examine how networks change with the removal of electrode contacts, and not upon the addition of contacts. For an intracranial EEG study that has parsimoniously captured seizure onset and spread, using our statistical sub-sampling method may miss critical electrode contacts and thus erroneously characterize the network as unreliable. However, for these patients that have very clear seizure onset and propagation, personalized network models of epilepsy may not be required to guide surgical planning. Additionally, we do not know how metrics would change under small deviations of electrode placement within the error bounds of appropriate surgical targeting. The jackknife analysis results varied somewhat across time, frequency band, and choice of seizure, reflecting the different states of the network. These observations underscore the fact that spatial sampling is one of several sources of bias of the network statistics. Further exploration of the sensitivity of network statistics to the choice of seizure, time, and frequency band will be necessary for useful clinical implementation. Also, additional steps toward translating this work into clinical care will require expanding our data set to include larger numbers of patients, stereo EEG implantations and those treated with focal laser ablation and perhaps brain stimulation devices.

While in this study we implemented only data-driven metrics that describe underlying network properties, there is significant interest in fitting generative models of neural population dynamics to brain networks. One such example, the Epileptor model (Jirsa et al. 2014), is a neural mass model that describes many relevant epileptic dynamics and is currently under study as a clinical trial in Europe to guide epilepsy surgery (Proix et al. 2018). Our network resampling approach may also be used for generative neurophysiologic models and give confidence to their clinical utility.

### 4.5 Conclusions

The field of network science provides a promising set of tools for understanding epilepsy dynamics and for surgical planning. However, the robustness of empirical estimates of network statistics to incomplete electrode sampling is not well understood. We have shown variability across network measures in robustness to incomplete sampling. Network measures are equally vulnerable to the removal of electrode contacts near versus distant from the seizure onset zone. Robustness to incomplete sampling is highly heterogeneous across patients, and jackknife resampling is a simple algorithm to obtain patient-specific confidence in the results of network statistics. The choice of individual network models should be based upon the intended application and on the clinical certainty that important seizure generators have been sampled by intracranial implants.

## Supplemental Materials

**Supplemental Figure 1:**
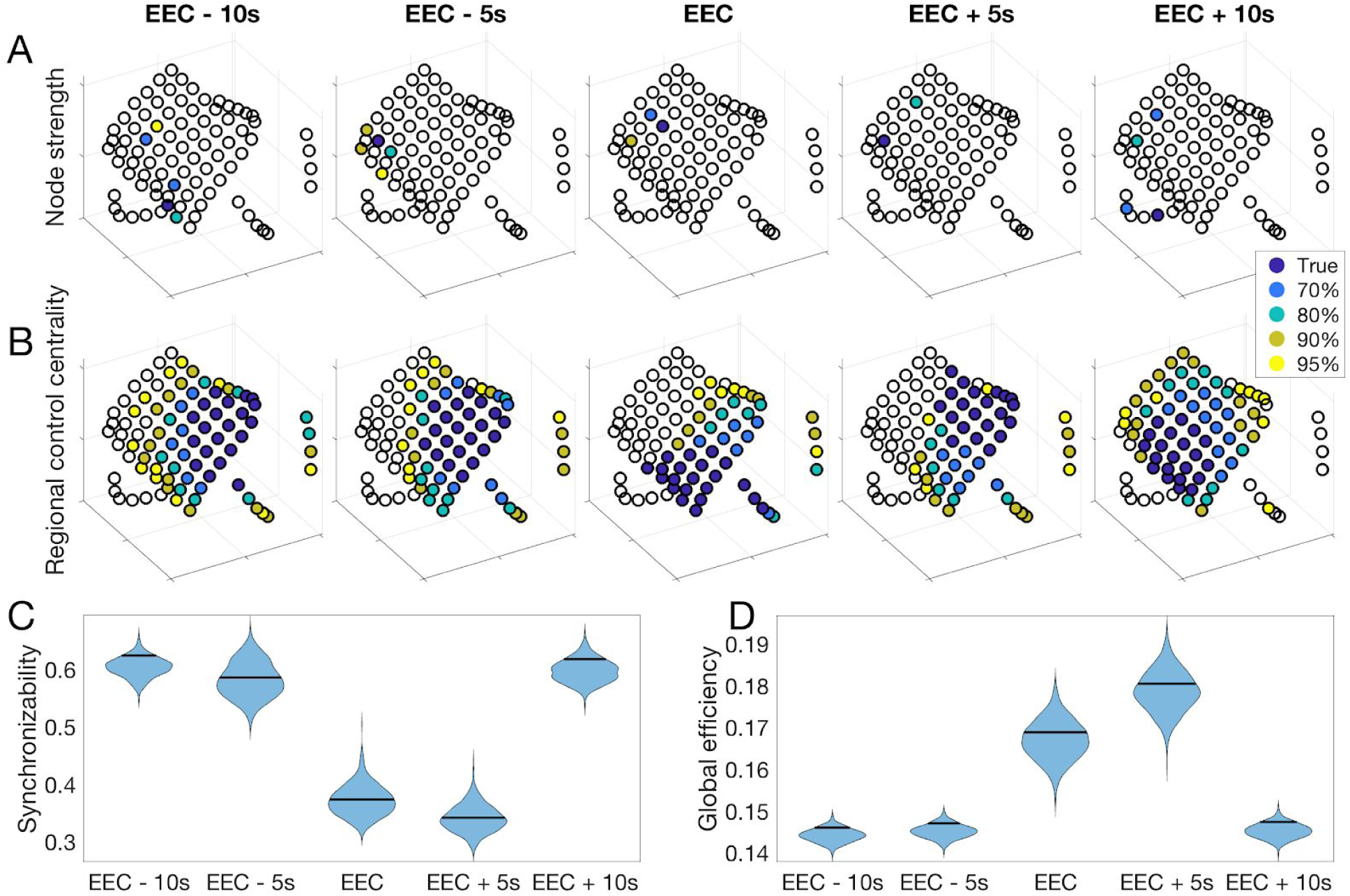
Time dependence of jackknife resampling results. All data shown is for a single example patient (HUP064). A: The location of the electrode with the highest node strength, as well as the electrodes accounting for various percentages of highest node strength occurrences in 1,000 random jackknife resampling networks at five different time periods. B: The location of the most synchronizing region, and the regions accounting for various percentages of the most synchronizing region occurrences in 1,000 random jackknife resampling networks at five different time periods. C: The distribution of synchronizability values obtained with 1,000 random jackknife resamplings for each time period. Horizontal lines denote synchronizability in the original non-resampled network. D: The distribution of global efficiency values obtained with 1,000 random resamplings for each time period. Horizontal lines denote global efficiency in the original non-resampled network.

**Supplemental Table 1.** Clinical and electrode information. 2-year post-surgical outcomes were based on the International League Against Epilepsy (ILAE) classification system (class 1-6). HUP: Hospital of the University of Pennsylvania, M: male, F: female, mo: months, LFL: left frontal lobe, LFPL: left frontoparietal lobe, LTL: left temporal lobe, RTL: right temporal lobe, RFTL: right frontotemporal lobe, RFL: right frontal lobe, MCD: malformation of cortical development; FCD: focal cortical dysplasia, HS/MTS: hippocampal sclerosis/mesial temporal sclerosis.

| Patient ID | Sex | Age at onset | Age at surgery | Localization | Pathology | ILAE outcome | Grids | Strips | Depths |
| --- | --- | --- | --- | --- | --- | --- | --- | --- | --- |
| HUP064 | M | 3 | 22 | LFL | MCD | 1 | 64 | 22 | 0 |
| HUP065 | M | 2 | 36 | RTL | MCD | 1 | 64 | 0 | 0 |
| HUP068 | F | 13 | 28 | RTL | HS/MTS | 1 | 63 | 20 | 0 |
| HUP070 | M | 12 | 33 | LFPL | MCD | 2 | 63 | 0 | 0 |
| HUP073 | M | 5 | 40 | RFL | MCD | 1 | 0 | 52 | 0 |
| HUP074 | F | 5 | 25 | LTL | MCD | 1 | 63 | 29 | 22 |
| HUP075 | F | 52 | 57 | LTL | HS/MTS | 5 | 58 | 34 | 14 |
| HUP078 | M | 2 mo | 54 | LTL | HS/MTS | 4 | 63 | 24 | 14 |
| HUP080 | F | 35 | 41 | LTL | Gliosis | 2 | 62 | 18 | 16 |
| HUP082 | F | 34 | 56 | RTL | HS/MTS | 1 | 64 | 14 | 8 |
| HUP083 | M | 8 | 29 | LPL | Gliosis | 2 | 59 | 22 | 0 |
| HUP086 | F | 18 | 25 | LTL | Gliosis | 1 | 60 | 32 | 8 |
| HUP087 | M | 19 | 24 | LFL | Tumor/Vascular<br>/Infection | 3 | 60 | 12 | 12 |
| HUP088 | M | 13 mo | 24 | LFL | HS/MTS | 1 | 0 | 43 | 11 |
| HUP094 | F | 20 | 48 | RTL | Gliosis | 2 | 0 | 64 | 20 |
| HUP105 | M | 27 | 39 | RTL | Tumor/Vascular<br>/Infection | 1 | 0 | 43 | 12 |
| HUP106 | F | 24 | 45 | LTL | HS/MTS | 2 | 64 | 36 | 16 |
| HUP107 | M | 5 | 36 | RTL | HS/MTS | 1 | 64 | 38 | 16 |
| HUP111A | F | 28 | 40 | RTL | HS/MTS | 1 | 0 | 32 | 16 |
| HUP111B | F | 28 | 40 | RTL | HS/MTS | 1 | 47 | 42 | 12 |
| HUP116 | F | 51 | 58 | RTL | unknown | 1 | 0 | 0 | 50 |
| Study012 | M | 23 | 37 | RFL | Gliosis | 1 | 57 | 22 | 0 |
| Study016 | F | 5 | 36 | RFTL | Gliosis | 4 | 47 | 14 | 0 |
| Study017 | M | unknown | unknown | RTL | unknown | 4 | 0 | 0 | 16 |
| Study019 | M | 31 | 33 | LTL | Gliosis | 5 | 60 | 28 | 8 |
| Study020 | M | 5 | 10 | RFL | Gliosis | 4 | 40 | 16 | 0 |
| Study022 | F | 11-20 | 21-30 | LTL | Gliosis | 5 | 42 | 12 | 0 |
| Study028 | M | 4 | 5 | LFPL | Gliosis | 4 | 64 | 5 | 0 |
| Study029 | F | unknown | unknown | RTL | Gliosis | 5 | 23 | 30 | 8 |

**Supplemental Table 2.**
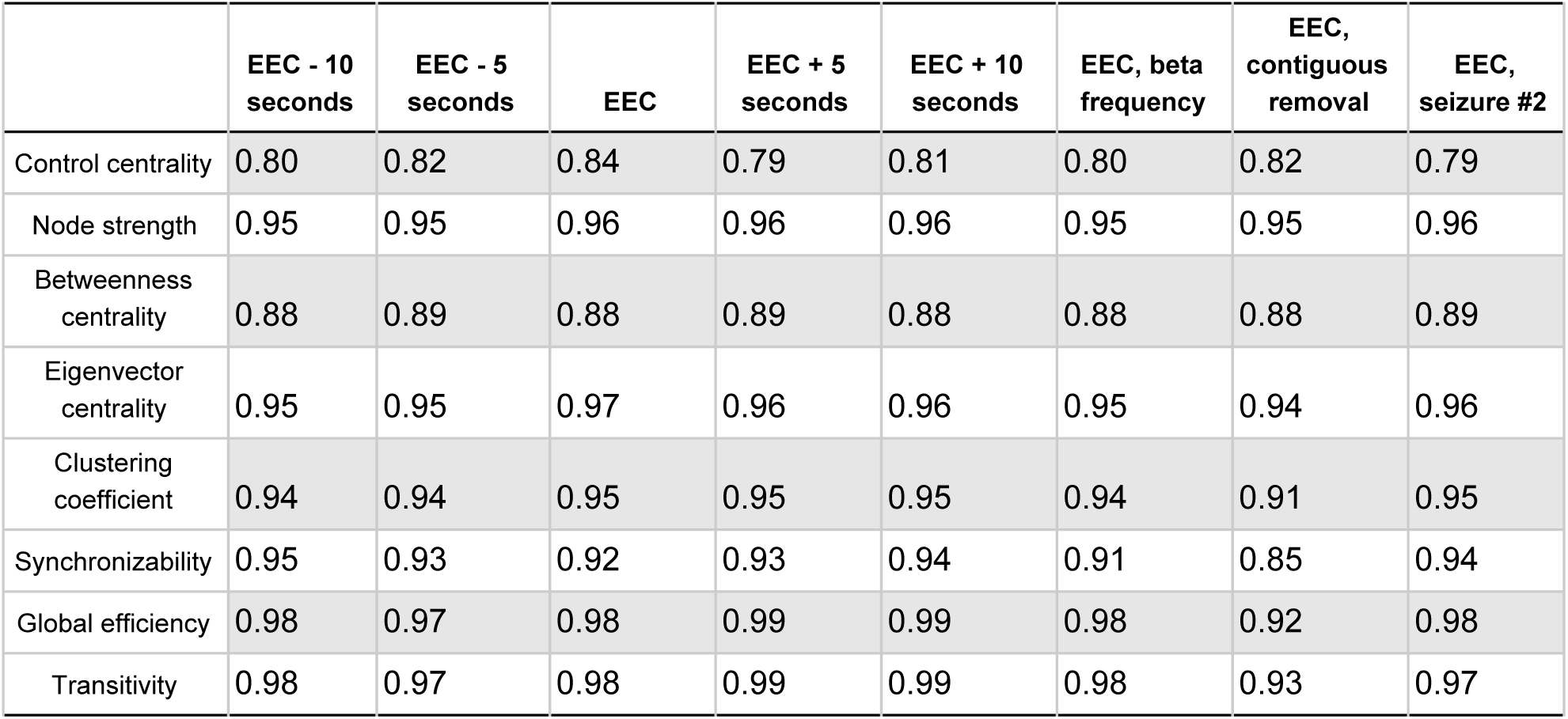
Metric reliability for alternative conditions. The metric reliability, defined in the text, for alternative conditions including different time periods relative to the earliest electrographic change (EEC) (as opposed to the primary analysis on the EEC), beta frequency coherence (as opposed to the primary analysis on high gamma frequency coherence), removal of a contiguous set of electrodes (as opposed to random set), and the second seizure (as opposed to the first seizure).

|  | EEC - 10<br>seconds | EEC - 5<br>seconds | EEC | EEC + 5<br>seconds | EEC + 10<br>seconds | EEC, beta<br>frequency | EEC,<br>contiguous<br>removal | EEC,<br>seizure #2 |
| --- | --- | --- | --- | --- | --- | --- | --- | --- |
| Control centrality | 0.80 | 0.82 | 0.84 | 0.79 | 0.81 | 0.80 | 0.82 | 0.79 |
| Node strength | 0.95 | 0.95 | 0.96 | 0.96 | 0.96 | 0.95 | 0.95 | 0.96 |
| Betweenness<br>centrality | 0.88 | 0.89 | 0.88 | 0.89 | 0.88 | 0.88 | 0.88 | 0.89 |
| Eigenvector<br>centrality | 0.95 | 0.95 | 0.97 | 0.96 | 0.96 | 0.95 | 0.94 | 0.96 |
| Clustering<br>coefficient | 0.94 | 0.94 | 0.95 | 0.95 | 0.95 | 0.94 | 0.91 | 0.95 |
| Synchronizability | 0.95 | 0.93 | 0.92 | 0.93 | 0.94 | 0.91 | 0.85 | 0.94 |
| Global efficiency | 0.98 | 0.97 | 0.98 | 0.99 | 0.99 | 0.98 | 0.92 | 0.98 |
| Transitivity | 0.98 | 0.97 | 0.98 | 0.99 | 0.99 | 0.98 | 0.93 | 0.97 |

**Supplemental Table 3.** Metric reliability for all removal percentages. The metric reliability, defined in the text, for all removal percentages tested. All data is shown for the time period at the EEC.

|  | EEC, 80% | EEC, 60% | EEC, 40% | EEC, 20% |
| --- | --- | --- | --- | --- |
| Control centrality | 0.62 | 0.70 | 0.76 | 0.84 |
| Node strength | 0.72 | 0.84 | 0.92 | 0.96 |
| Betweenness<br>centrality | 0.62 | 0.70 | 0.79 | 0.88 |
| Eigenvector<br>centrality | 0.75 | 0.86 | 0.92 | 0.97 |
| Clustering<br>coefficient | 0.60 | 0.78 | 0.88 | 0.95 |
| Synchronizability | 0.65 | 0.77 | 0.85 | 0.92 |
| Global efficiency | 0.82 | 0.91 | 0.95 | 0.98 |
| Transitivity | 0.84 | 0.91 | 0.95 | 0.98 |

**Supplemental Table 4.** Association between metric agreement and distance of removed electrodes from seizure onset zone for alternative conditions. Values denote the *t*-statistic evaluating the patient-aggregated Fisher’s transformed Spearman rank correlations for the distance-agreement associations. The method for calculating the agreement-distance association is described in the text. Positive values indicate that the metric is more sensitive to removing electrodes near the resection zone. Values are shown for different time periods relative to the earliest electrographic change (EEC) (as opposed to the primary analysis on the EEC), beta frequency coherence (as opposed to the primary analysis on high gamma frequency coherence), and the second seizure (as opposed to the first seizure). No association was significant for α = 0.05/8 (Bonferroni correction).

|  | EEC - 10<br>seconds | EEC - 5<br>seconds | EEC | EEC + 5<br>seconds | EEC + 10<br>seconds | EEC, beta<br>frequency | EEC,<br>seizure #2 |
| --- | --- | --- | --- | --- | --- | --- | --- |
| Control centrality | -0.09 | 0.02 | 0.80 | 2.64 | 1.15 | 0.49 | 1.06 |
| Node strength | 1.47 | 0.07 | 1.25 | 1.30 | -0.83 | 1.76 | 0.95 |
| Betweenness<br>centrality | -0.83 | -0.22 | -0.95 | -1.22 | 0.13 | 0.04 | -0.31 |
| Eigenvector<br>centrality | 1.60 | 0.22 | 1.02 | 1.72 | -0.47 | 1.92 | 1.37 |
| Clustering<br>coefficient | 1.75 | 0.02 | 1.23 | 1.53 | -0.84 | 1.16 | 1.22 |
| Synchronizability | -0.30 | 0.80 | -0.26 | 0.85 | 0.84 | -0.11 | -1.06 |
| Global efficiency | 2.00 | 0.79 | 0.74 | 0.54 | 2.31 | 1.67 | -0.61 |
| Transitivity | -0.02 | 1.43 | 0.37 | 0.30 | 1.70 | 0.55 | -1.39 |

**Supplemental Table 5.** Association between metric agreement and distance of removed electrodes from seizure onset zone for alternative removal percentages. Values denote the *t*-statistic evaluating the patient-aggregated Fisher’s transformed Spearman rank correlations for the distance-agreement associations. The method for calculating the agreement-distance association is described in the text. Positive values indicate that the metric is more sensitive to removing electrodes near the resection zone. Values are shown for different percentages of removed electrodes. All results are for the time period at the earliest electrographic change (EEC), high gamma frequency coherence, and the first seizure. No association was significant for α = 0.05/8 (Bonferroni correction).

|  | EEC, 80% | EEC, 60% | EEC, 40% | EEC, 20% |
| --- | --- | --- | --- | --- |
| Control centrality | -0.58 | 0.61 | 2.23 | 0.80 |
| Node strength | 0.42 | 1.53 | 1.53 | 1.25 |
| Betweenness<br>centrality | 0.74 | 0.16 | 0.49 | -0.95 |
| Eigenvector<br>centrality | 0.47 | 1.72 | 1.46 | 1.02 |
| Clustering<br>coefficient | 0.55 | 1.46 | 1.44 | 1.23 |
| Synchronizability | 1.28 | 2.25 | 0.95 | -0.26 |
| Global efficiency | 0.34 | 0.89 | 0.90 | 0.74 |
| Transitivity | 0.54 | 0.54 | 0.55 | 0.37 |

**Supplemental Table 6.** Results of jackknife resampling method for alternative conditions. Each nodal value shows the average number of electrodes accounting for 95% of all occurrences of the maximum metric value (minimum for control centrality and regional control centrality) across 1,000 jackknife resamples. Each global value shows the width of the 95% jackknife confidence interval of the network metric across 1,000 jackknife resamples. Values shown are for alternative conditions including different time periods relative to the earliest electrographic change (EEC) (as opposed to the primary analysis on the EEC), beta frequency coherence (as opposed to the primary analysis on high gamma frequency coherence), removal of a contiguous set of electrodes (as opposed to random set), and the second seizure (as opposed to the first seizure).

|  | EEC - 10<br>seconds | EEC - 5<br>seconds | EEC | EEC + 5<br>seconds | EEC + 10<br>seconds | EEC, beta<br>frequency | EEC,<br>contiguous<br>removal | EEC,<br>seizure #2 |
| --- | --- | --- | --- | --- | --- | --- | --- | --- |
| Node strength | 3 | 3 | 3 | 3 | 3 | 4 | 3 | 3 |
| Betweenness<br>centrality | 4 | 4 | 4 | 5 | 5 | 5 | 4 | 4 |
| Eigenvector<br>centrality | 3 | 4 | 3 | 3 | 3 | 4 | 3 | 3 |
| Clustering<br>coefficient | 3 | 3 | 3 | 3 | 3 | 4 | 3 | 3 |
| Control centrality | 9.5 | 11 | 9 | 8 | 7 | 10 | 6 | 9 |
| Regional control<br>centrality | 52 | 51 | 48 | 51 | 47.5 | 52 | 50 | 44 |
| Synchronizability | 0.09 | 0.10 | 0.09 | 0.09 | 0.09 | 0.09 | 0.13 | 0.09 |
| Global efficiency | 0.01 | 0.02 | 0.02 | 0.02 | 0.02 | 0.01 | 0.02 | 0.02 |
| Transitivity | 0.01 | 0.01 | 0.01 | 0.01 | 0.01 | 0.01 | 0.02 | 0.01 |

